# GDF5 modulation of MuSC pool as a potential therapeutic benefit for DMD

**DOI:** 10.1101/2025.10.14.682261

**Authors:** Christel Gentil, Aly Bourguiba, Amélie Vergnol, Bruno Cadot, Zoheir Guesmia, Lucile Saillard, Pierre Meunier, Bénédicte Hoareau-Coudert, Sestina Falcone, Lorenzo Giordani, France Pietri-Rouxel

## Abstract

Duchenne muscular dystrophy (DMD) is a fatal disease caused by dystrophin deficiency, leading to degeneration of the entire musculature. To improve muscle pathophysiology and gene therapy for DMD, we investigated the potential of growth differentiation factor 5 (GDF5) in the DMD mdx mouse model. We showed that the overexpression of GDF5 in the muscle improved its histology, reduced inflammation, modulated regeneration and induced the appearance of *de novo* fibers. We demonstrated that muscle satellite cells (MuSCs) are targeted by GDF5 which enhanced their proliferation and slowed down their myogenic commitment and finally their fusion. When combined with AAV-mediated microdystrophin gene therapy, the leading therapeutic strategy, GDF5 further increased the number of microdystrophin-positive fibers compared to gene therapy alone. These findings highlight GDF5 as a promising modulator of DMD pathology and provide the first evidence of a synergistic effect of the combination of GDF5-based intervention and AAV-microdystrophin treatment.

## Introduction

Duchenne muscular dystrophy (DMD) is a devastating muscular disorder caused by the lack of the dystrophin protein. As dystrophin is essential for the integrity of muscle fibers, its absence results in muscle necrosis along with cyclic degeneration and regeneration. Initially, regeneration in DMD disease is supported by the proliferation and differentiation of muscle satellite cells (MuSCs). However, regenerative potential of these cells becomes progressively exhausted, rendering muscle repair inefficient and leading to muscular dysfunction^1^. In addition, recent studies revealed defective MuSCs polarity and reduced number of myocytes in mdx muscle following acute injury^2,3^. Nevertheless, the preservation of muscle renewing capacity depends on a complex cellular network. Indeed, even though MuSCs sustain the formation of new myofibers in response to injuries, other elements of the niche (for example immune cells and interstitial progenitors) have been shown to participate to muscle maintenance and regeneration^4,5^.

Among the therapeutic strategies developed for Duchenne muscular dystrophy (DMD), clinical trials based on microdystrophin (µDystrophin) gene transfer mediated by Adeno Associated Virus (AAV) administration (AAV-µDys) are underway and show encouraging results with improvement or a stabilization of motor functions and a reduction in blood CPK enzyme levels. However, there are challenges facing such approaches, including pre-existing anti-capsid immunity, anti-transgene immunity, serious cardiac events, the functionality of µdystrophin, and the long-term transgene persistence^6–8^. Indeed, these strategies target the degenerating muscle, so their long-term success highly depends on maintaining muscle mass for as long as possible and is thus heavily in need of preserving muscle fiber integrity, decreasing regeneration process and thus, limiting muscle wasting. With the aim of improving the pathophysiology of DMD disease and thus optimizing gene therapy treatments, such as µDystrophin supplementation, we focused on muscle trophic factors. In terms of muscle tissue homeostasis, components of the Bone Morphogenetic Protein (BMP) family that bind BMP receptor 1 (BMPR1) have been shown to induce muscle hypertrophy via SMAD (fusion of *Caenorhabditis elegans Sma* genes and the *Drosophila Mad*, or mothers against decapentaplegic) 1/5/8 phosphorylation and Akt/mTOR pathway activation^9^. BMP signaling is also active in adult regenerating muscle, in which it regulates MuSCs proliferation/differentiation^10,11^. Noteworthy, downregulation of BMPs target genes, such as the Inhibitor of differentiation (Id) factors involved in the control of cell cycle and differentiation^12^, delays muscle regeneration by altering MuSC-dependent myogenesis^13^. In addition, BMP signaling has been shown as possible modulator of other resident cells present in the muscle niche and, indeed, BMPR1 activation inhibits fibro/adipogenesis during muscle repair^14^.

Muscle atrophy and chronic inflammation are hallmarks contributing to the pathophysiology of different muscle disorders, such as age-related sarcopenia^15^, congenital myopathies^16^, myositis^17^ or DMD^18^. We have been pioneers in discovering that overexpression of the BMP14 protein, also called Growth Differentiation Factor 5 (GDF5), was able to prevent muscle mass loss, to decrease inflammation and increase of anti-oxidant-related markers’ expression and to maintain force generation in aged mice^19,20^. Despite the evidence that GDF5 is a positive modulator of muscle mass homeostasis and participates in the regulation of muscle repair, its potential applications to preserve muscle quality in disease such as DMD has not been explored.

Here, we demonstrated that GDF5 overexpression (GDF5-OE) in muscle has a beneficial effect on DMD pathophysiology by reducing necrotic and inflammatory areas, modulating regeneration process and, importantly, by inducing hyperplasia in mdx mice. Single nuclei RNA sequencing (snRNA-seq) allowed identification of MuSCs as preferential targets of GDF5 which modulates the balance between quiescence and activation states of these cells. Furthermore, we confirmed *in vitro* that GDF5 increased the pool of mdx-derived MuSCs by delaying the progression of these cells toward myogenic differentiation, a delay due to the inhibition of the fusion process. Going further in the goal of gene therapy improvement for DMD, we combined GDF5 overexpression with AAV-µDys administration and revealed a synergistic effect with the appearance of higher number of µDys positive small fibers compared to the AAV-µDys condition alone.

Overall, we identified a novel role of GDF5 in preserving dystrophic muscle integrity and assessed the therapeutic potential of combining GDF5 with AAV-µDys to reduce the dystrophic features and enhance the benefits of gene therapy.

## Results

### GDF5 overexpression reduces necrosis and inflammation hallmarks

To evaluate the impact of GDF5-OE in a dystrophic model, we used mdx mice, a well characterized DMD model, whose muscles undergo a temporary major crisis with myofiber degeneration followed by regenerative process between 3 and 8 weeks of age^21^. The effect of GDF5 OE was investigated by injecting 1 week old-mdx *Tibialis Anterior* (TA) with an AAV2/9 carrying the GDF5 transgene, (mdx-GDF5) or with an AAV2/9-control containing backbone pSMD2 without transgene sequence (mdx). Muscles were collected at 2- and 8-weeks post-injection (2 p.i. and 8 p.i., respectively) from mouse euthanasia at 3 (before the crisis) and 9 weeks (after the crisis) of age and compared with muscles from age-matched wild type (WT) mice (Fig 1a). First, we validated the overexpression of the GDF5 transcript and protein in mdx-GDF5 up to 2 p.i. (Fig 1b-c) and confirmed that the GDF5 pathway was activated as demonstrated by the increase of the SMAD1/5 complex phosphorylation (p-SMAD1/5) and of its nuclear accumulation (Fig S1a-b) as well as of the *id4* transcript, a gene known to be induced by GDF5^12^ (Fig S1c).

**Figure 1.**
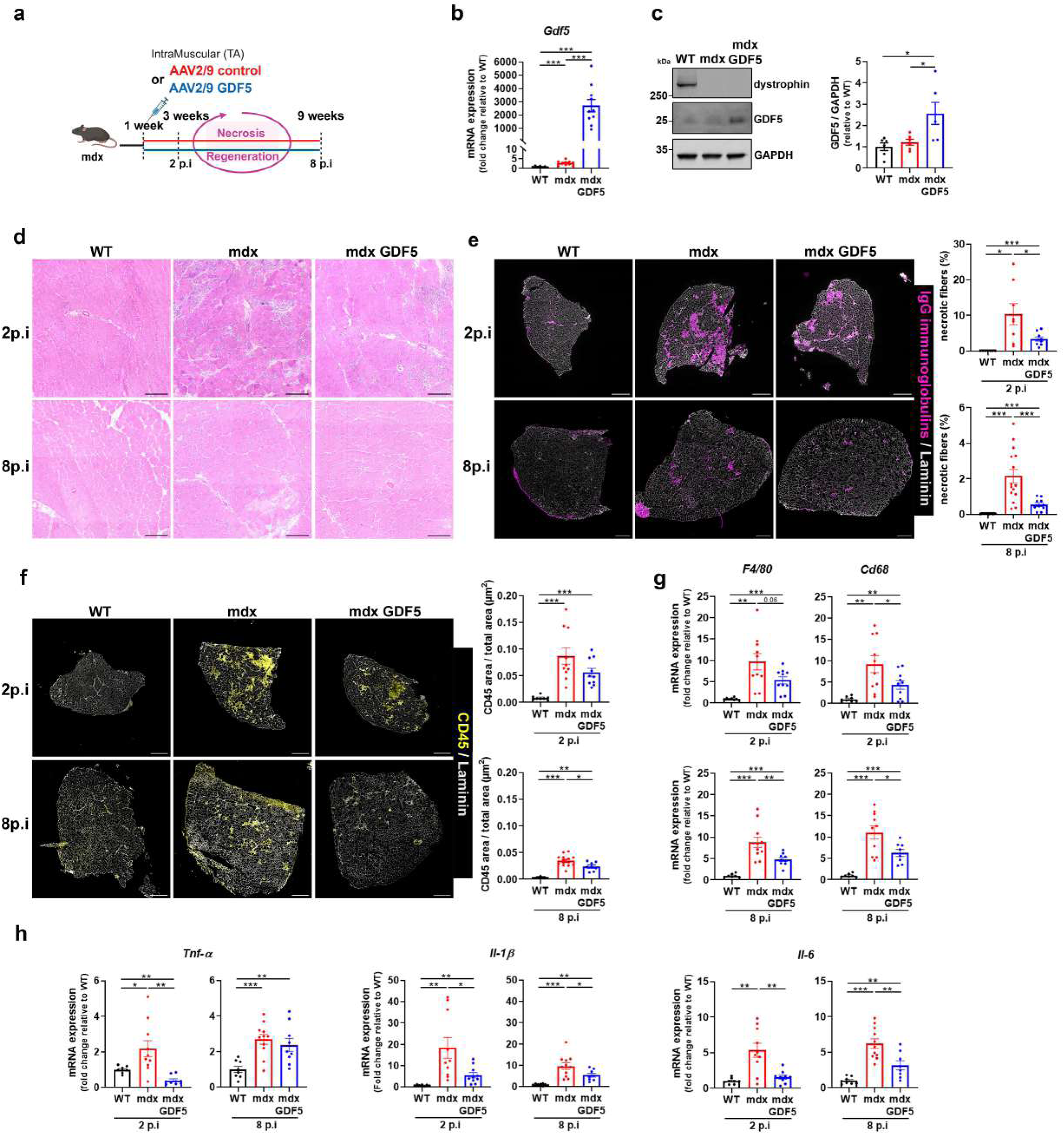
GDF5 overexpression reduces necrosis and inflammation hallmarks. **a**: Schematic *in vitro* experimental design. 1-week-old mdx were injected into *Tibialis Anterior* (TA) with AAV2/9-control (mdx) or with AAV2/9-GDF5 (mdx-GDF5). Molecular and functional analyses were performed in TAs 2 and 8 weeks post-injection (2- and 8-p.i.) and compared with age-matched wild-type (WT). **b**: RT-qPCR of *Gdf5* expression in TA from WT, mdx and mdx-GDF5, 2 p.i. (n=7-10). **c**: Representative Western blot of TA lysates from WT, mdx and mdx-GDF5, 2 p.i., probed for dystrophin, GDF5, and GAPDH. GDF5 levels was quantified and normalized to GAPDH (n=6). **d**: Representative images of TA cryosections from WT, mdx and mdx-GDF5, 2- and 8-p.i., stained with hematoxylin and eosin, scale bar 250 μm. **e-f**: Representative images of TA cryosections from WT, mdx and mdx-GDF5, 2- and 8-p.i. (e) Cryosections stained with IgG immunoglobulins highlighting necrotic fibers (purple) and laminin (grey), scale bar 500 μm. Quantification of necrotic fibers relative to total fibers. (2p.i.: n=7-9) (8p.i.: n=8-15). (f) Cryosections stained with CD45 highlighting inflammatory cells (yellow) and laminin (grey), scale bar 500 μm. Quantification of CD45 area relative to total area (2p.i.: n=8-10) (8p.i.: n=3-14). **g-h**: RT-qPCR to measure (g) *F4/80* and *Cd68* expression and (h) *Tnf-α, Il-1β and Il-6* expression in TA from WT, mdx and mdx-GDF5, 2 and 8p.i. (2p.i.: n=8-10).(8p.i.: n=7-11). Data are presented as means ± s.e.m. *P-values* were calculated by Brown One-Way Forsythe ANOVA test followed by Unpaired t with Welch’s correction (b, c, e, f, g, h); Ordinary One-Way ANOVA test followed by Uncorrected Fisher LSD. (f, h).

To determine whether GDF5-OE might exert beneficial effect on the dystrophic phenotype of mdx mice, TA sections were examined after hematoxylin/eosin staining. Histological analysis revealed reduced dystrophic features and infiltration area (Fig 1d) as well as necrotic fibers’ ratio, quantified by immunoglobulin G uptake^22^ in both, 2 p.i. and 8 p.i. muscles (Fig 1e). In addition, we observed a reduced inflammatory cell infiltration in treated TA of mdx at 8 p.i., marked by the decrease of area positive for Leucocyte common antigen CD45 and by the reduce of the expression of transcripts of the macrophage markers *Cd68* and *F4/80*^23^ (Fig 1f-g). We confirmed the effect of GDF5-OE on inflammatory status by measuring the expression of the well-known pro-inflammatory cytokines *Tnf-α*^24^, *Il1-β*^25^, *Il-6*^26^. Their expression was decreased in 2 p.i. mdx-GDF5 muscles for all markers, and this diminution was maintained for *Il1-β* and *Il-6* compared to mdx 8 p.i. TAs (Fig 1h).

Altogether, these data demonstrate a benefit of GDF5-OE in mdx muscle with an improvement of DMD muscle histology associated with a reduction of the necrotic fiber area and a decline of inflammation.

### GDF5 overexpression leads to muscle hyperplasia

Muscle regeneration was then investigated by the quantification of centrally nucleated fibers. A decrease of their percentage in 8 p.i. mdx-GDF5 compared to mdx muscles was observed, suggesting a modulation of this process by GDF5-OE (Fig 2a, b). As GDF5 has been described as a key factor promoting muscle mass maintenance during aging^20^, we weighed the mdx TAs in the different conditions. At 2 p.i., while mdx muscles displayed atrophy compared to WT, likely due to necrosis, muscle mass was no longer significantly different from that of WT following GDF5 OE. At 8 p.i., untreated mdx muscles exhibited a pathological increase in muscle weight probably due to fibrosis and/or cellular infiltration as previously described in mdx TA muscles^2^. No significant difference in muscle mass was observed between GDF5-treated and untreated mdx muscles (Fig. S2a). However, at this time of treatment, histological analysis revealed hyperplasia with an increase of the total number of fibers in mdx-GDF5 mainly due to increased number of small myofibers (<2000 µm^2^) associated with a decrease of large fibers’ number (<3000 µm^2^) which usually marks pathological hypertrophy^27^ (Fig 2c, d). Modification of fiber type distribution is a hallmark of dystrophic muscle and has been reported to be modified in mdx compared to WT TA^28^. Our results showed that hyperplasia due to GDF5-OE was accompanied by a shift of fiber type in 8 p.i. mdx-GDF5 compared to mdx muscles. Indeed, we observed an increase of the slow-twitch oxidative types I and of the fast-twitch oxidative-glycolytic type IIX while the fast-twitch glycolytic IIB fibers were decreased (Fig S2b). Notably, oxidative-glycolytic fibers type IIX display higher fatigue resistance in routine continuous activities such as posture maintenance and walking^29^ and in DMD pathological context fast muscle fibers are preferentially affected^30^. We next quantified muscle renewing by Embryonic Myosin Heavy Chain (eMyHC) staining^31^. We found a reduced number of labelled fibers in 2 p.i. mdx-GDF5, suggesting a delayed in the onset of the regeneration process. In 8 p.i. mdx-GDF5, eMyHc positive fibers percentage was similar to that mdx muscle but the labelled fibers were smaller (Fig 2e-g) suggesting a delay in fiber maturation likely reflecting a slower initiation of regeneration at 2 p.i.

**Figure 2.**
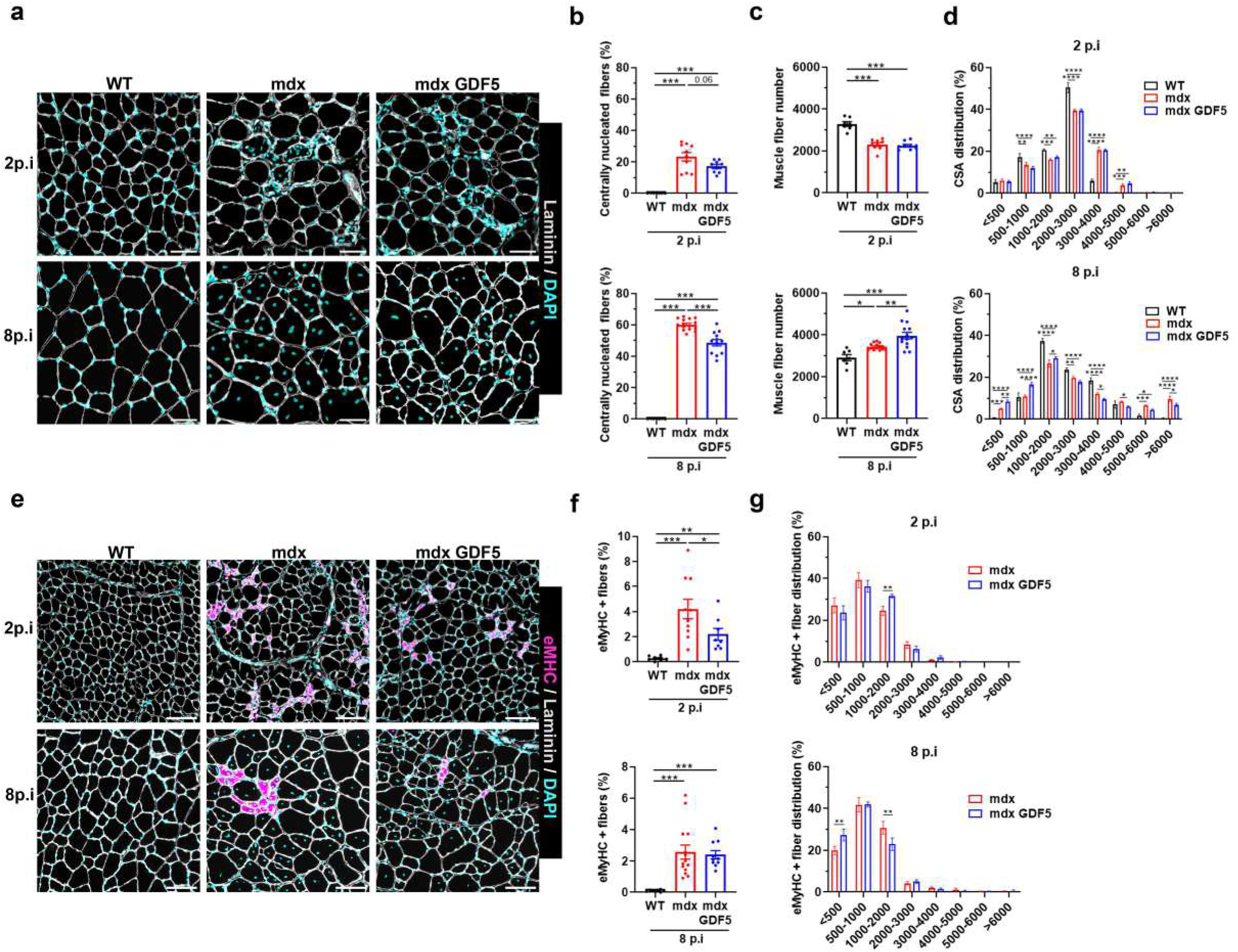
GDF5 overexpression leads to muscle hyperplasia. **a-d**: (a) Representative images of TA cryosections from WT, mdx and mdx-GDF5, 2- and 8-p.i., immunostained with laminin (grey) and DAPI (cyan), scale bar 50 μm. (b) Percentage of centrally nucleated fibers among total fibers (2p.i.: n=7-10) (8p.i.: n=7-16). (c) Quantification of muscle fiber number in whole TA (2p.i.: n=7-9) (8p.i.: n=7-14). (d) Distribution of fibers according to cross section area. (2p.i.: n=7-10) (8p.i.: n=7-14). **e-g**: (e) Representative images of TA cryosections from WT, mdx and mdx-GDF5, 2- and 8-p.i., immunostained with embryonic MHC (eMHC) (purple), laminin (grey) and DAPI (cyan), scale bar: 100µm. (f): Percentage of eMHC positive fibers among total fibers (2p.i.: n=8-10) (8p.i.: n=7-14) (g) Distribution of eMHC positive fibers according to cross section area (2p.i.: n=9-10) (8p.i.: n=10-14). Data are presented as means ± s.e.m. *P-values* were calculated by Brown Forsythe One-Way ANOVA test followed by Unpaired t with Welch’s correction (b, c, f); Ordinary One-Way ANOVA test followed by Uncorrected Fisher LSD. (c); Two-Way ANOVA test followed by Uncorrected Fisher LSD comparisons test (d, g).

In summary, mdx muscles overexpressing GDF5 during 8 weeks exhibited a modulation of regeneration characterized by both hyperplasia marked by the emergence of newly formed fibers and a delayed maturation of fibers accompanied by a shift of fiber type.

### GDF5 overexpression drives changes in MuSCs dynamics in dystrophic muscle

To obtain an overview of the different cell populations expressing GDF5 after muscle AAV-GDF5 treatment, a snRNA-seq was performed on TAs from WT, mdx, and mdx-GDF5 mice at 2- and 8 -p.i. (Fig 3a). Clusters were annotated based on canonical markers, allowing us to identify all major known populations consistent with previous reports. In addition, we detected a population of assembling fibers (Flnc⁺) and a population of Meg3⁺ fibers, similar to those described by Petrany et al. at postnatal day 21 in the TA of healthy mice (Fig 3b-c)^10,32,33^. GDF5 expression was validated according to mouse groups, time post-injection and cell populations (Fig 3c). In 2 p.i. mdx-GDF5 muscles, data revealed its transcript was predominantly in muscle fibers and MuSCs (Fig 3d, Fig S3a). After 8 weeks of treatment, we observed a significant decrease of the GDF5 transcript expression (P=3.6e-4) in all cell populations compared to the 2 p.i. condition, even though it remained present at low level (Fig 3d, Fig S3a). To correlate GDF5 transcript expression to the presence of AAV, we quantified viral genome (vg) content in treated muscles. Surprisingly, the vg content was found higher in 2 p.i. mdx-GDF5 compared to mdx muscles suggesting improved maintenance of viral genomes within the muscle fibers. Indeed, in dystrophic muscle, fiber degeneration leads to the loss of viral genomes, a process that appears to be attenuated in muscles treated with GDF5 (Fig S3b). However, in 8 p.i. mdx muscles, treated with ether AAV-control or AAV-GDF5, the AAV vg content was significantly decreased compared to 2 p.i. mdx muscles revealing the loss of the vg overtime as already described^8,34^.

**Figure 3.**
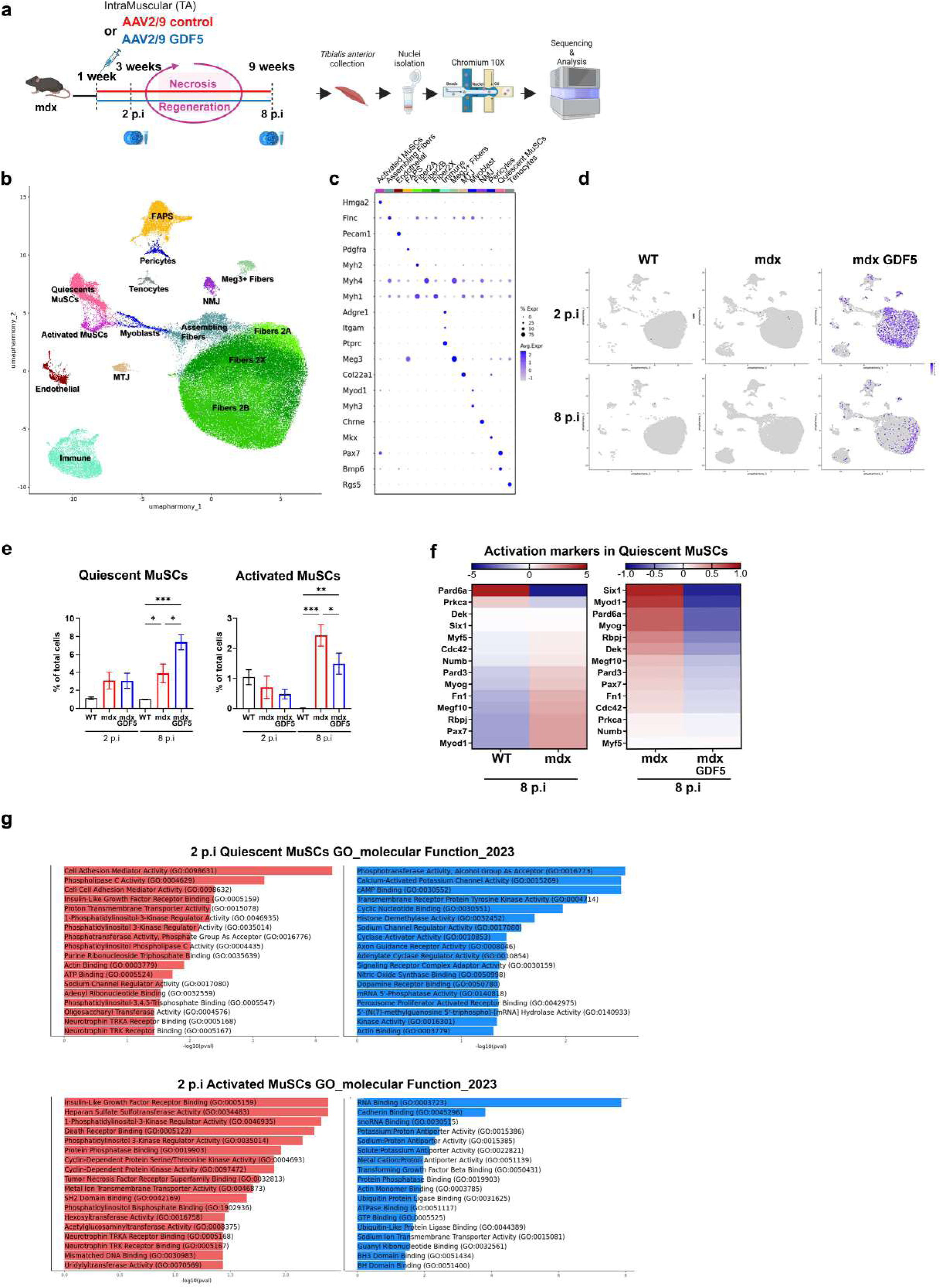
GDF5 overexpression drives changes in MuSCs dynamics in dystrophic muscle. **a**: Schematic Single nucleus RNA sequencing experimental design. 1-week-old mdx were injected into *Tibialis Anterior* (TA) with AAV2/9-control (mdx) or with AAV2/9-GDF5 (mdx-GDF5). Nuclei isolation was performed following the Chromium Nuclei Isolation Kit (10x Genomics) 2- and 8-p.i. Nuclei were encapsulated in barcoded beads and then sequenced. **b**: Unbiased clustering of snRNA-seq data from 2- and 8-p.i. TAs represented on a UMAP plot. Each major cell type was represented in a different color after clusterization. **c**: Dotplot showing marker genes for major cell types in TA. **d**: Feature plots assessing *Gdf5* expression in WT mdx and mdx-GDF5 2- and 8-p.i. *p-values* were calculated with a negative binomial distribution adjusted by Benjamini-Hochberg rank test^65^ **e**: Percentage of quiescent muscle stem cells (left) and activated muscle stem cells (MuSCs) (right) in WT, mdx and mdx-GDF5 TA 2- and 8-p.i. *P* values were calculated by mixed-effects analysis followed by Fisher LSD test. **f**: Heatmap of Log2 fold change gene expression of activation markers between WT and mdx (left) or mdx and mdx-GDF5 (right) quiescent MuScs 8 p.i. **g**: Top 18 upregulated (red) and downregulated (blue) Gene Ontology (GO) terms 2 p.i. in mdx-GDF5 compared to mdx in Quiescents MuSCs (up) and Activated MuSCs (down).

To further investigate the impact of GDF5-OE on muscle homeostasis in mdx, and given that MuSCs play a pivotal role in maintaining muscle integrity^35^, we assessed their activation status. We first measured the proportion of quiescent MuSCs and found it to be higher in mdx-GDF5 muscles at 8 p.i. compared to both mdx and WT mice (Fig. 3e). In contrast, the proportion of activated MuSCs was significantly lower in mdx-GDF5 than in mdx muscles, although still higher than in WT mice. At 8 p.i., analysis of gene expression in quiescent MuSCs from mdx muscles revealed elevated expression of several activation associated markers as *MyoD*, *MyoG*, *Fn1*, *Numb*, *Dek* and *Megf10*^36^ while their expression was reduced in quiescent MuSCs from mdx-GDF5 muscles (Fig 3f). These findings suggest that GDF5-OE could modulate the activation state of MuSCs in mdx muscles.

Both the differences between mdx and WT, and those between mdx and mdx-GDF5 observed at 8 p.i. can be traced back to molecular events already initiated at 2 p.i.. To explore the molecular basis of these events and their impact we performed Gene Ontology (GO) term analysis on both quiescent and activated MuSCs from these muscles, comparing mdx to age-matched WT controls (Fig S3c) and mdx-GDF5 to age-matched mdx controls (Fig. 3g). The analysis revealed upregulation of cell adhesion pathways in quiescent mdx-GDF5 MuSCs and increased cyclin-dependent kinase activity in activated MuSCs, suggesting changes in proliferation and/or differentiation signalling. Notably, one of the most downregulated GO terms was “cadherin binding,” a process implicated in the differentiation of neural stem cells^37^.

To investigate GDF5-induced changes in cell-cell communication, we performed CellChat analysis comparing mdx and mdx-GDF5 at 2 p.i. (Fig S3d). This revealed differences in the NCAM and VCAM pathways. NCAM signalling was increased in both quiescent and activated MuSCs from mdx-GDF5 relative to mdx control muscles (Fig S3e). Notably, VCAM signalling, undetectable in mdx MuSCs, were restored between immune cells and quiescents MuSCs in mdx-GDF5 muscle (Fig S3f) as VCAM interactions are known to support MuSCs proliferation and survival, this suggests that GDF5-OE enhances cell-cell communication within MuSCs niche.

Altogether, these data indicate that AAV-GDF5 treatment mainly influences MuSCs fate. Specifically, GDF5-OE led to an increase of the proportion of quiescent MuSCs and a decrease of activated MuSCs. Additionally, our analysis revealed that AAV-GDF5 treatment significantly impacted MuSC cell adhesion pathways with an upregulation of the expression of their markers observed in quiescent MuSCs. In particular, we measured increased NCAM and VCAM pathways as well as upregulation of cyclin-dependent kinase activity and downregulation of cadherin binding, indicating that GDF5 might promote proliferation and inhibit differentiation of these cells.

### GDF5 overexpression modulates myogenic regulatory factors and increases the pool of MUSCs

These results led us to further investigate how GDF5-OE might influence MuSC myogenic properties beyond gene expression profiles. In particular, we focused on the shift observed in DMD MuSCs from quiescent and activated states toward more committed myogenic progenitor states. First, we quantified Pax7-positive cells, a hallmark of MuSCs, involved in early myogenic commitment^39^. Quantification of Pax7-positive nuclei revealed a significant higher number of MuSCs in muscles upon GDF5-OE at 2 p.i. compared to WT and mdx muscles (Fig 4a). Additionally, double immunostaining for Pax7 and the proliferation marker Ki67^40^ showed an increased proportion of PAX7/KI67 positive nuclei in mdx-GDF5 muscles compared to WT and non-treated mdx, indicating a higher proportion of proliferating MuSCs that had not yet committed to differentiation (Fig 4b).

**Figure 4.**
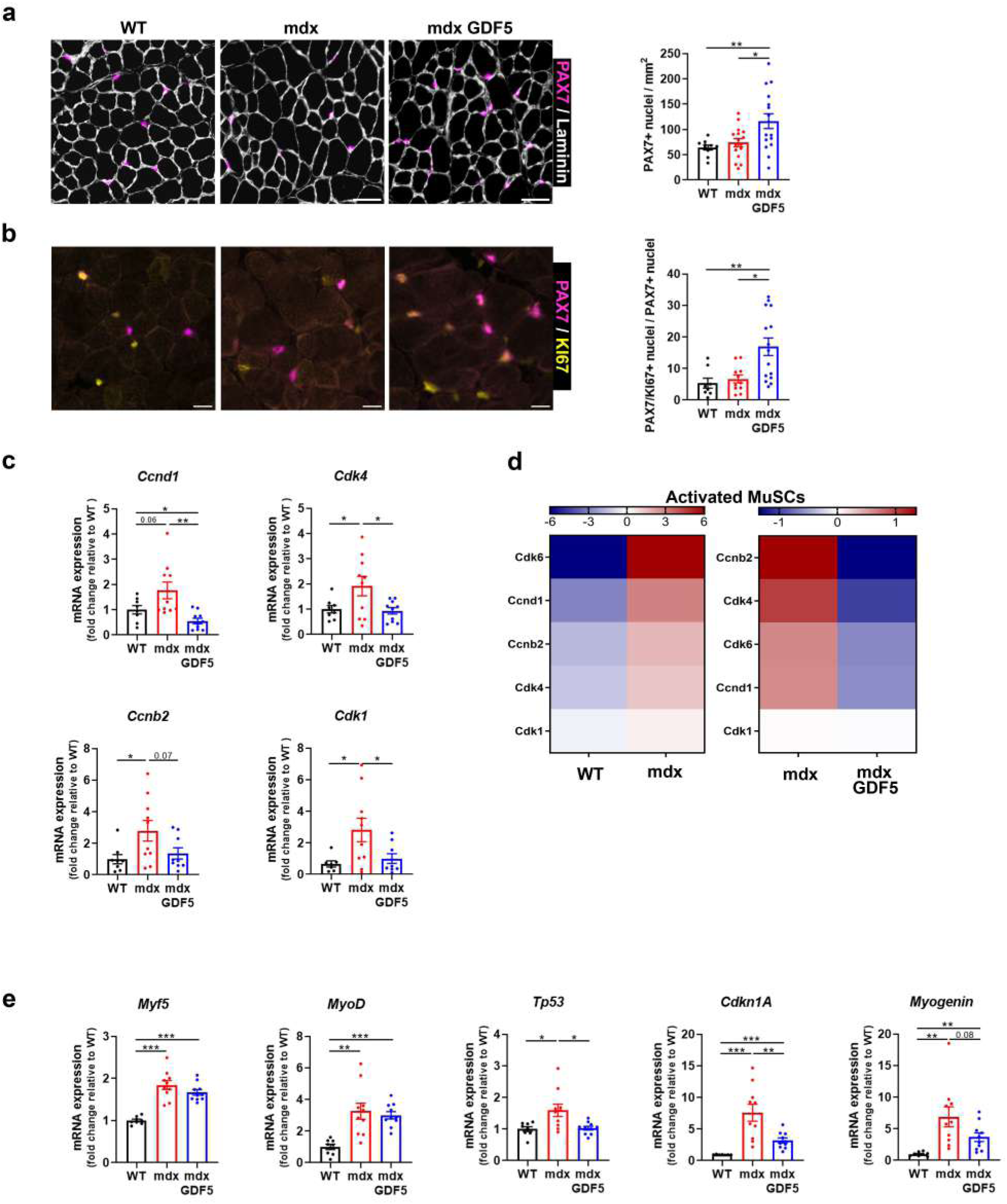
GDF5 overexpression modulates myogenic regulatory factors and increases the pool of MUSCs. **a-b**: (a) Representative images of TA cryosections from WT, mdx and mdx-GDF5 (2 p.i.). (b) Cryosections stained with PAX7 (purple) and laminin (grey), scale bar: 50µm. Percentage of PAX7 positive nuclei per mm^2^. (n=10-16). (b) Cryosections stained with PAX7 (purple) and KI67 (yellow). Scale bar: 20µm. Percentage of PAX7/KI67 positive nuclei relative to PAX7 positive nuclei. (n=8-15). **c**: RT-qPCR of *Ccnd1*, *Cdk4*, *Ccnb2 and Cdk1* expression in TA from WT, mdx and mdx-GDF5 2 p.i. (n=7-10). **d**: Heatmap of Log2 fold change gene expression of cyclin and cdk between WT and mdx (left) or mdx and mdx-GDF5 (right) activated MuScs 2 p.i. **e**: RT-qPCR of Myf5, *MyoD*, Tp53, Cdkn1A and *Myogenin*, expression in TA from WT, mdx and mdx-GDF5 2 p.i. (n=8-10). Data are presented as means ± s.e.m. *p-values* were calculated by Brown Forsythe One-Way ANOVA test followed by Unpaired t with Welch’s correction (a, c, d); Ordinary One-Way ANOVA test followed by Uncorrected Fisher LSD (d); Kruskal-Wallis test followed by Uncorrected Dunn’s test (b).

Given the increased proportion of proliferating MuSCs in mdx-GDF5 muscles, we then examined the expression of key regulators of cell cycle progression. Cyclins and their cyclin-dependent kinase (CDK) partners orchestrate cell cycle progression and are essential for controlling MuSC proliferation and the transition between quiescent and activated states. In particular, cyclin D1, in association with CDK4, regulates the G1/S transition, while cyclin B2, together with CDK1, control the G2/M transition^41^.

The transcriptional expression of these cyclins was increased in mdx muscles compared to WT muscles, suggesting an abnormal proliferative response in the context of dystrophic muscle regeneration and repair, and was normalized in mdx-GDF5 muscles (Fig. 4c).

Furthermore, analysis of snRNA-seq data revealed that upregulated cyclin pattern was characteristic of activated mdx MuSCs and that was decreased after treatment with GDF5 (Fig. 4d). To determine whether such proliferative abnormalities in mdx muscle were associated with dysregulation of myogenic transcription factor expression^2^, an alteration that has previously been reported, and to assess whether GDF5-OE could modulate this process, we analyzed the expression of myogenic commitment markers (Fig. 4e). We found that the expression of *MyoD* and *Myf5* was increased in mdx muscle and was not affected by GDF5 OE. However, *myogenin*, which controls the expression of genes involved in cell cycle exit and myoblast fusion into multinucleated myotubes^42,43^, was upregulated in mdx and tended to be reduced by GDF5-OE. Finally, p21 and p53, which together halt the cell cycle allowing MuSCs to exit proliferation and initiate muscle differentiation^44,45^, were overexpressed in mdx muscles^3^ and reduced in mdx following GDF5 treatment.

These data demonstrate that GDF5-OE leads to a normalisation of the quiescent/activate state of MuSCs, downregulated myogenic commitment transcript expression suggesting that it could maintain myoblasts in a proliferative state, preserving their capacity to expand before committing to differentiation.

### GDF5 overexpression modulates proliferation and differentiation of MuSCs

To better understand the impact of GDF5 on MuSCs, this population was isolated from WT and 2 p.i. mdx muscles treated or not with GDF5 by fluorescence-activated cell sorting (FACS) using the profile CD31-, CD45-, Sca1-, and VCAM+^46^ (Fig 5a, 5b). Consistently with what was observed *in vivo*, the number of MuSCs per milligram isolated from fresh muscle was higher in mdx-GDF5 compared to mdx (Fig 5c).

**Figure 5.**
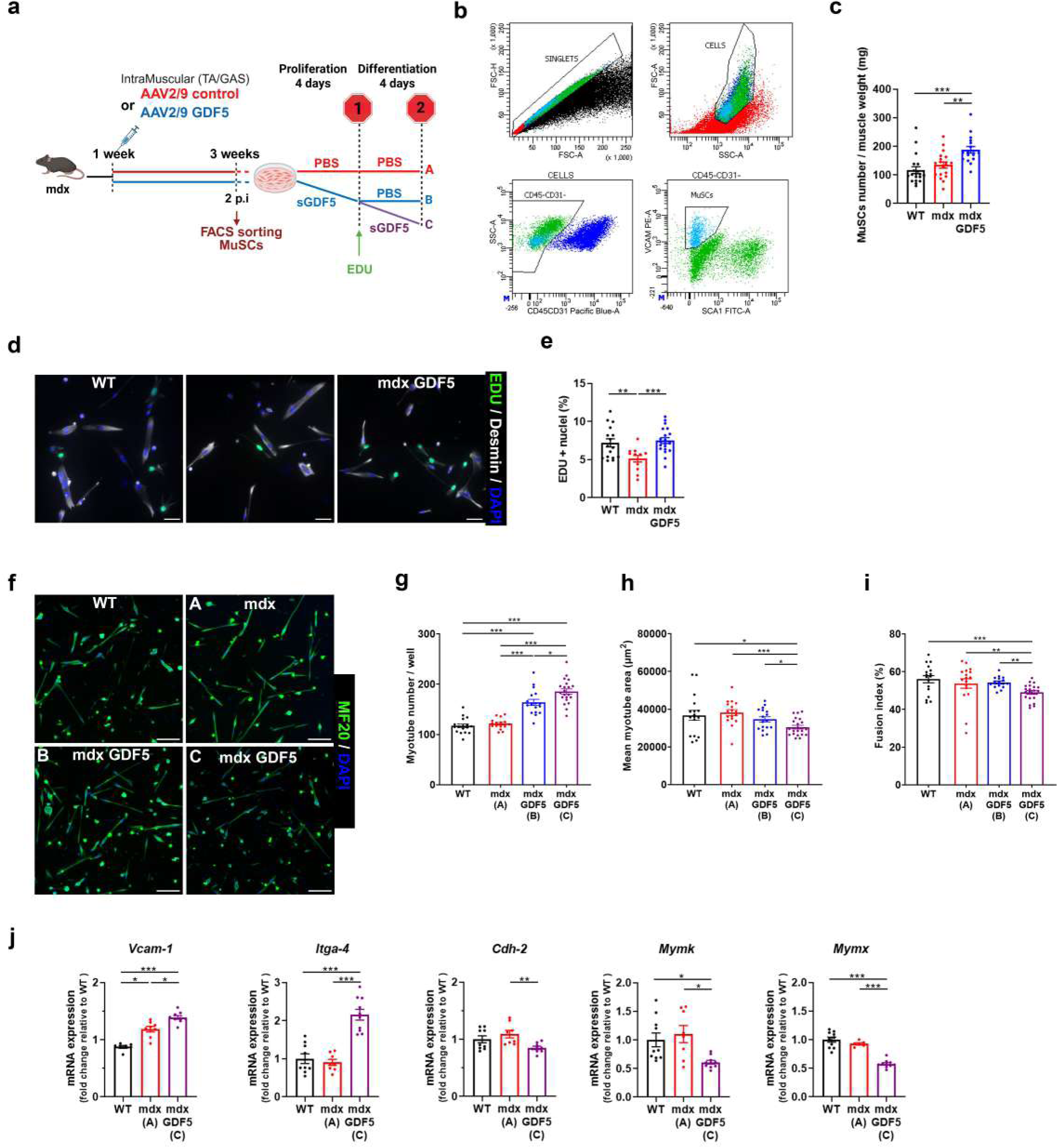
GDF5 overexpression modulates proliferation and differentiation of MuSCs. **a-b**: Schematic of *in vitro* experimental design. 1-week-old mdx were injected into *Tibialis Anterior* (TA) and *Gastrocnemius* (GAS) with AAV2/9-control (mdx) or with AAV2/9-GDF5 (mdx-GDF5) 2 p.i., muscle satellite cells (MuSCs) were isolated from mononuclear cells by FACS using the profile CD31⁻/CD45⁻/Sca1⁻/VCAM⁺ and compared to WT. MuSCs were cultured for 4 days in proliferation medium with or without sGDF5 (STOP1-panels d, e) or for 4 days in proliferation medium with or without sGDF5 followed by 4 days in differentiation medium with or without sGDF5 (STOP2-panels f-j). **c**: Quantification of MuSC number of muscles (TA and GAS) from WT, mdx and mdx-GDF5, 2 p.i., after FACS sorting relative to muscle weight (mg). (n=23-25) **d**: MuSC proliferation rate assessed by EdU incorporation 4 hours before the fixation. (d) Representative image of proliferating satellite cells immunostained with EDU (green), DAPI (blue) and Desmin (grey), scale bar 50µm. (e) Percentage of EDU positive nuclei from the total number of nuclei. (n=16-20). **f**: Representative image of differentiating cells immunostained with MF20 (green) and DAPI (blue). Scale bar 500µm. **g-i**: (g) Quantification of the number of myotubes per well. (h) Mean myotubes area. (i) Fusion index. (n=17-21). **j**: RT-qPCR of *Vcam-1, Itga-4, Cdh-2, Mymk* and *Mymx* expression in MuSCs from WT or mdx, untreated or treated during proliferation and differentiation with sGDF5. (n=7-15). Data are presented as means ± s.e.m. *p-values* were calculated by Brown Forsythe One-Way ANOVA test followed by Unpaired t with Welch’s correction (g, h, j); Ordinary One-Way ANOVA test followed by Uncorrected Fisher LSD (c, e, j); Kruskal-Wallis test followed by Uncorrected Dunn’s test (I, j).

The sorted MuSCs were then cultured in proliferation medium with or without synthetic GDF5 (sGDF5) protein^20^ for four days (Fig 5a - STOP1) or for 4 days in proliferation medium with or without sGDF5 followed by 4 days in differentiation medium with or without sGDF5 (Fig 5a - STOP2).

*In vitro* experiments using 5-Ethynyl-2’-deoxyuridine (EdU) reagent, incorporated into the DNA of dividing cells (Fig 5a), revealed that sGDF5 treatment normalized the percentage of EdU-positive nuclei, suggesting that treatment with GDF5 might help to maintain myoblasts in a proliferating state and prevents premature differentiation (Fig5d-e). To evaluate the effect of GDF5 on myoblast differentiation, MuSCs were exposed to sGDF5 during proliferation and either maintained or withdrawn during differentiation (Fig. 5a). We found that treatment with sGDF5 during proliferation alone increased the number of myoblasts, ultimately leading to a higher number of myotubes at the end of differentiation (Fig. 5f-g). When sGDF5 was maintained during differentiation, the number of myotubes exceeded that observed under proliferation-only conditions. However, both the fusion index and the mean surface area of the myotubes were reduced (Fig. 5h-i). It should be noted that continuous treatment with sGDF5 throughout proliferation and differentiation appeared to maintain the cells at the expense of terminal differentiation.

Myotube formation depends not only on the expression of myogenic transcription factors, but also on the expression of fusion and adhesion related transmembrane proteins which play an essential role in regulating stem cell quiescence, activation and differentiation^47^. Considering our data from snRNA sequencing analysis showing the effect of GDF5 on VCAM signalling, and knowing that VCAM-1 and integrin α4 (Itga-4) promote MuSCs expansion and prevent premature progression to a more differentiated state^38^, we assessed the transcript levels of these two factors in the MuSCs cultured with sGDF5 during both proliferation and differentiation (Fig 5a, Fig 5j). Consistent with previous findings, maintaining GDF5 treatment throughout differentiation increased expression of *VCAM-1* and *Itga-4*, suggesting that myoblasts were retained in a proliferative state. In contrast, we observed downregulation of *Cdh2* expression, which mediates cell-cell adhesion, as well as that of *Mymk* and *Mymx* which coordinate the membrane fusion process essential for myotube formation^48–50^.

Collectively, these results suggest that sGDF5 treatment could expand the pool of MuSCs probably by limiting their ability to fully differentiate, which correlates with the delayed myogenic commitment observed in mdx-GDF5 muscles.

### Combination therapy using GDF5 OE and AAV-µdystrophin

Given that GDF5-OE promoted muscle hyperplasia in dystrophic TA, likely by inducing expansion of the MuSCs pool and slowing early differentiation, we hypothesized that it could potentiate the efficacy of AAV-μDys gene therapy. To test this hypothesis, we performed a sequential injection protocol, a first injection of AAV2/9-GDF5 to expand the MuSCs reservoir, followed by an injection of AAV2/10 carrying the μDys transgene (*mdx*-GDF5-μDys mice) (Fig. 6a). Importantly, ELISA analysis confirmed that prior administration of AAV2/9 did not induce a humoral immune response that could have compromised subsequent AAV2/10 delivery through the generation of cross-reactive neutralizing antibodies (Fig. S4). As expected, RT-qPCR analyses confirmed high expression of *Gdf5* transcripts in *mdx*-GDF5-μDys muscles (Fig. 6b), and *μDys* expression was verified both at the transcript level (Fig. 6c) and protein level via Western blot (Fig. 6d).

**Figure 6.**
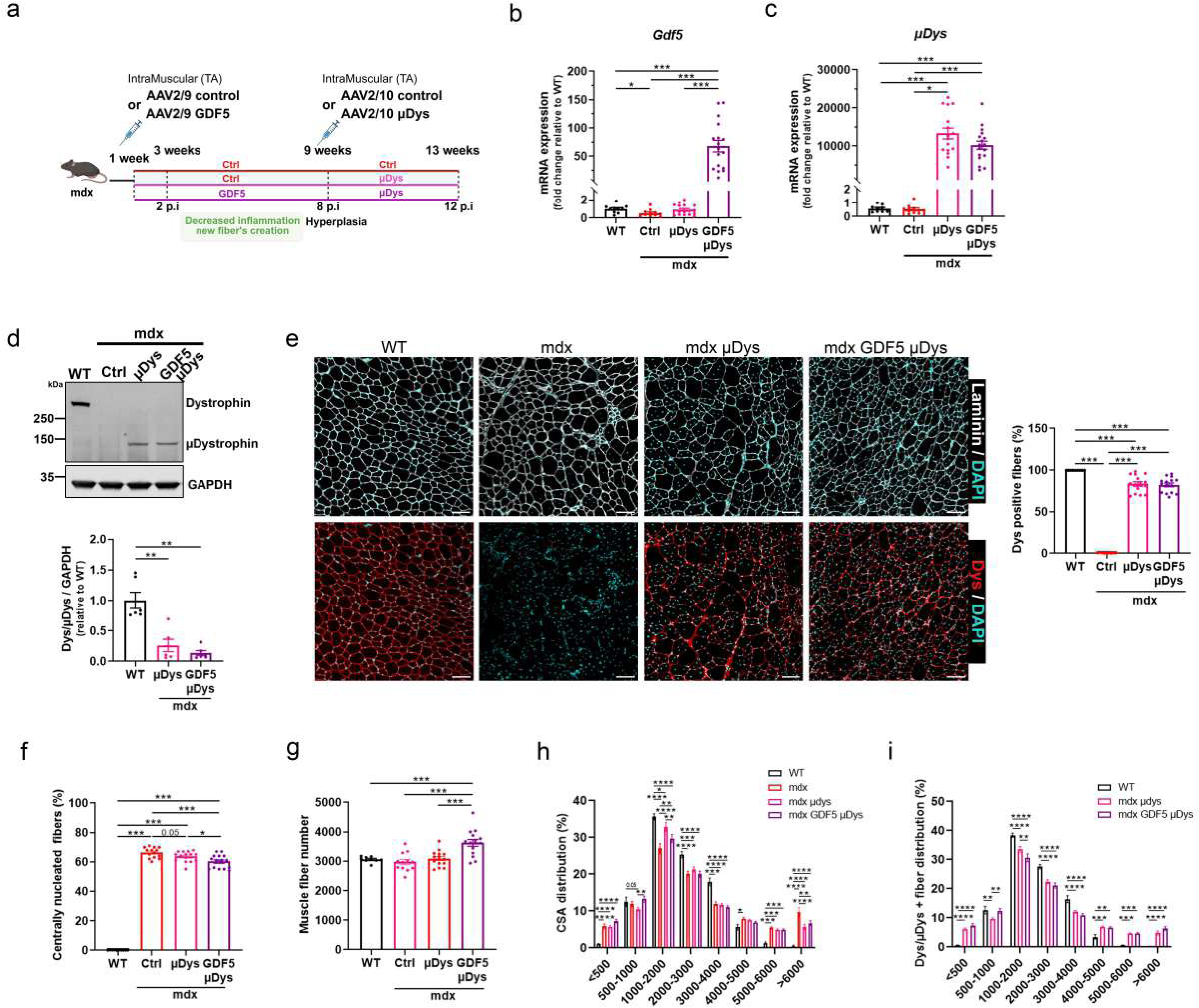
Combination therapy using GDF5-OE and AAV-µdystrophin. **a**: Schematic *in vivo* experimental design. 1-week-old mdx were injected into TA with AAV2/9-control or with AAV2/9-GDF5 and then 8 weeks later, the mice received a second injection of AAV2/10- µ-Dystrophin (mdx-µDys or mdx-GDF5-µDys). These treated mice were compared to mdx mice treated with AAV2/9-control and then with AAV2/10-control (mdx-Ctrl). The analysis was performed 4 weeks later and compared with age-matched wild-type (WT). **b-c**: RT-qPCR of (b) *Gdf5* and (c) *µdystrophin* expression in TA from WT, mdx-Ctrl, mdx-µDys and mdx-GDF5-µDys (n=9-17). **d**: Representative Western blot of TA lysates from WT, mdx-Ctrl, mdx-µDys and mdx-GDF5-µDys. Blots were probed for dystrophin, µdystrophin, and GAPDH; Dys and µDys levels quantified and normalized to GAPDH (n=6). **e**: Representative images of TA cryosections from WT, mdx-Ctrl, mdx-µDys and mdx-GDF5-µDys, immunostained with laminin (grey), dystrophin or µdystrophin (red) and DAPI (cyan), scale bar 100 μm. Quantification of dystrophin positive fiber among total fibers. (n=9-15). **f**: Percentage of centrally nucleated fibers among total fibers. (n=10-15). **g**: Quantification of muscle fiber number in whole TA. (n=9-15). **h**: Distribution of fibers according to cross section area. (n=10-15). **i**: Distribution of Dys positive fibers according to cross section area. (n=10-15). Data are presented as means ± s.e.m. *p-values* were calculated by Brown Forsythe One-Way ANOVA test followed by Unpaired t with Welch’s correction (b,c,e,f,g), Kruskal-Wallis test followed by Uncorrected Dunn’s test (d); Ordinary One-Way ANOVA test followed by Uncorrected Fisher LSD (f); Two-Way ANOVA test followed by Uncorrected Fisher LSD (h,i).

The assessment of the number of µDys-positive fibers revealed no significant difference between *mdx*- μDys and *mdx*-GDF5-μDys mice (Fig. 6e). However, muscles from the group receiving the combined treatment exhibited a significantly reduced proportion of centrally nucleated fibers compared to the μDys-only group suggesting decreased degeneration/regeneration cycles leading to an improved muscle morphology (Fig. 6f). Quantification of the total number of fibers revealed their significant increase in the *mdx*-GDF5-μDys group, accompanied by a higher percentage of small-diameter fibers (<1000 μm²) compared to the μDys group (Fig. 6g, h). Furthermore, analysis of the size distribution of dystrophin-positive fibers showed a notable enrichment of small μDys-expressing myofibers (1000 μm²) in the combined treatment group (Fig. 6i) indicating that the newly formed fibers induced by GDF5 OE were efficiently transduced by the AAV-μDys vector.

Collectively, these findings support the evidence of a synergistic effect of GDF5-OE and μDys gene transfer, enhancing its therapeutic impact by expanding the population of μDys-expressing fibers and potentially improving the permanence and efficacy of the treatment.

## Discussion

DMD is a progressive degenerative disease in which preservation of muscle integrity is critical for maximizing the efficacy of gene therapies. In this study, we investigated the potential benefit of GDF5 overexpression in the mdx mouse model and demonstrated that GDF5 could attenuate key pathophysiological features of DMD by reducing necrosis/regeneration cycles. In particular, we demonstrated that GDF5 acts by promoting the normalization of the MuSCs proliferation/differentiation balance, which is disrupted in dystrophic muscles. MuSCs from GDF5-treated mice, were maintained in a proliferative state contributing to the formation of new fibers that were efficiently transduced by AAV-µDys. This expansion of µDys-positive fibers represents a significant enhancement in the potential efficacy of gene replacement therapy.

The mdx mouse model was selected owing to its well-characterized phase of intense phase of necrosis-regeneration between 3 and 9 weeks of age, allowing us the study of GDF5 effects on these processes over a short time frame. The therapeutic potential of GDF5 was therefore tested by inducing its overexpression through AAV9-mediated gene delivery into the TAs of one-week-old mdx mice, and the experiments were performed just before the onset of the acute crisis and after complete regeneration. At both time points, GDF5-OE conferred clear benefits, including histological improvements characterized by reduced necrosis, decreased inflammation, and protection against atrophy. These results are consistent with prior reports describing GDF5 as a regulator of muscle mass preservation during aging^20^ and in delaying the onset of necrosis intervertebral disc degeneration^51^.

We further attributed the beneficial effects of GDF5 to its capacity to regulate muscle remodelling by influencing the balance between MuSC quiescence and activation. In DMD muscles, absence of dystrophin induces MuSC dysfunction in the form of as loss of polarity^52–54^ leading to exacerbated differentiation of these cells^2^.

By limiting aberrant myogenic commitment, GDF5 preserved MuSC quiescence, maintaining the stem cell reservoir. Complementary *in vitro* experiments confirmed that sGDF5 treatment enhanced MuSC proliferation while delaying terminal differentiation by inhibiting cell fusion. Although this delay of differentiation process could appear inconsistent with the hyperplasia observed in mdx-GDF5 muscles at 8 weeks post-injection, transcriptomic analyses revealed a reduction in GDF5 expression at this time point, likely due to AAV genome loss in dystrophic tissue as previously described^8,34^. Consequently, previously expanded MuSC pools were released from GDF5-mediated proliferation, entered differentiation, and contributed to the emergence of new fibers, leading to hyperplasia and a shift toward oxidative, fatigue-resistant fiber type.

These findings align with studies showing that BMP signaling promotes MuSC proliferation by suppressing premature differentiation^11^. Interestingly, previous study examined the role of BMP signalling in regulating satellite cell function showing that its activation suppresses satellite cell differentiation^55^. More precisely, *Hatazawa et al*. described that the binding of GDF5 to its receptors on satellite cells inhibits their differentiation^56^.

While GDF5 delays MuSC differentiation to sustain their proliferative capacity, the molecular mechanisms underlying this effect required further investigation. With this objective, we analysed Gene ontology data and found that pathways involving VCAM and NCAM were among the most significantly modulated in mdx-GDF5 muscles 2 p.i.. Supporting these results, *in vitro* studies confirmed upregulation of *Vcam1* and *Itga4* in sGDF5-treated MuSCs compared to untreated WT or mdx MuSCs. It should be noted that the cell-cell adhesion protein VCAM-1 was described as allowing MuSCs to interact each other and with immune cells through α4β1 integrin, the highest-affinity VCAM-1 receptor, in an injured muscle niche to promote expansion of these cells and regulate their properties^38^.

The hyperplasia, observed in mdx 8 p.i. muscles, holds particular therapeutic relevance in the context of AAV-mediated µDys gene therapy. Combining GDF5-OE with AAV-µDys produced synergistic effects evidenced by an increased number of small µDys-positive fibers and improved muscle architecture. This synergy is likely driven by GDF5’s capacity to delay aberrant MuSC differentiation, thereby facilitating gene transfer efficiency and ensuring the persistence of newly generated fibers. Moreover, this combined approach directly addresses a key challenge in DMD therapy: the progressive depletion of MuSCs, which undermines long-term regenerative capacity and gene therapy durability. Looking ahead, we aim to replace AAV-mediated GDF5 delivery with systemic administration of synthetic GDF5 protein. This strategy would enable temporal control of treatment, allowing GDF5 exposure to be discontinued once an adequate MuSC pool is established, thereby preserving progenitor cells for later engagement during AAV-µDys therapeutic intervention. Beyond DMD, GDF5 may also benefit other myopathies marked by impaired regeneration and muscle wasting, such as congenital and limb-girdle muscular dystrophies or inflammatory myopathies. By sustaining MuSC proliferation, limiting premature differentiation, and protecting against atrophy, GDF5 could help preserving muscle homeostasis and complement gene- or cell-based therapies across a broad spectrum of neuromuscular disorders.

## Methods

### Animals

All animal procedures were reviewed and approved by an external ethics committee and the French Ministry of Higher Education and Scientific Research (Project authorization # 29280). Experiments were conducted on C57BL/10ScSn-Dmdmdx/J and C57BL/6 mice. The animals were housed under SPF conditions under 12h/12h light/dark conditions, with *ad libitum* access to food and water and cared for following the Directive 2010/63/EU.

### Plasmids and Adeno Associated Virus (AAV) production

pSMD2-GDF5 have been generated by direct cloning of GDF5 ORF (NM_008109.2), flanked by EcoRI and NheI sites (GeneArt string; ThermoFisher Scientific), in pSMD2 AAV2/9 under CMV promoter. pSMD2-control have been generated by direct cloning of pSMD2 without transgene sequence and pSMD2-md1 (microdystrophin) by direct cloning of pAAVSpc512JCMurine [George Dickson: Royal Holloway - University of London (RHUL)], both under spc5-12 promoter. AAV2/9 and AAV2/10 pseudotyped vectors have been prepared by the AAV production facility of the Center of Research in Myology, by transfection in 293 cells as described previously^19^. The final viral preparations were kept in phosphate-buffered saline (PBS) solution at −80°C. The particle titre (number of viral genomes per ml) was determined by quantitative PCR.

### *In vivo* treatments

For experiments on 1-week-old mice, animals were first anesthetized using isoflurane (4% induction, 3% maintenance) and then injected intramuscularly with AAV2/9-control or AAV2/9-GDF5 at 6.10^10^ viral genomes per ml (vg/ml) (10µl/ *Tibialis anterior* (TA) and 15µl/ *Gastrocnemius* (GAS). For experiments on 9-week-old mice, animals were first anesthetized using isoflurane (3% induction, 2% maintenance) and then injected intramuscularly with AAV2/10-control or AAV2/10-md1 at 6.10^11^ vg/ml (40µl/TA). At the end of procedure, mice were euthanized by cervical dislocation and TA and GAS muscles were then collected. TAs were frozen in liquid nitrogen–cooled isopentane for histological analysis and RNA extraction or TA and GAS muscles were used for satellite cells isolation.

### Fluorescence-Activated Cell Sorting (FACS)

For FACS, we used TA and GAS from mice injected at 7 days of age and treated for 2 weeks with AAV2/9-control or AAV2/9-GDF5. Muscles were isolated, minced and incubated first in muscle digestion buffer with 700-800 U/mL Collagenase II (Worthington) for 60 min à 37°C and then, with 1000U/ml Collagenase II (Worthington) and 11U/ml Dispase I (Life Technologies), for 30 min at 37° C and purified by filtration using 40 µm cell strainers. Satellite cell population was isolated from mononuclear cells using labelling extracellular antigens using APC anti-mouse CD31, APC anti-mouse CD45, Pacific Blue anti-mouse Ly-6A/E (Sca1) and PE anti mouse CD106 (VCAM1). Antibodies are listed in table S2.

### Cell cultures

After isolation, cells were cultured on plates pre-coated with Matrigel® (growth factor reduced, Corning). They were grown in DMEM (ThermoFisher Scientific) supplemented with 20% Fetal Bovine Serum (FBS) and gentamycin (50µg/ml, ThermoFisher Scientific), and then induced to differentiate using myogenic medium composed of DMEM with 2% Horse serum (HS) and gentamycin (50µg/ml). Cells were treated with or without synthetic GDF5 (100ng/ml, X’prochem) during both the proliferation and differentiation.

### EDU incorporation and detection

5-ethynyl-2′-deoxyuridine (EDU) was added in proliferating cells (final concentration of 10µM) 4 hours before fixation in 4% paraformaldehyde (PFA-Sigma-Aldrich). Cells were then incubated with the reaction solution (Click-iT EdU Cell Proliferation Assays, Thermofisher Scientific) for 30 minutes at room temperature to detect EDU prior to proceeding with the immunostaining protocol.

### Histology and Immunofluorescence

Muscle sections (10 μm thick) were performed on a cryostat (Leica Biosystems), fixed on glass slides and stored at -80°C.

For H&E staining, sections were fixed in 4% PFA for 10 min, washed in PBS and then stained in haematoxylin (Sigma-Aldrich) for 2 min and eosin (Sigma-Aldrich) for 30 sec. The muscle sections were further dried in gradually increasing concentration of ethanol/water solutions and, after fixation in 100% xylene, were mounted in Vectamount (Vector Laboratories).

For immunofluorescence procedures, cryosections (fixed or not with PFA 4%) or MuSCs (fixed with PFA 4%) for 10 min, were permeabilized with 0.5% Triton X-100 (Sigma-Aldrich) and blocked in PBS/4% bovine serum albumin (Sigma-Aldrich)/0.1% Triton X-100 for 1 h. Sections or cells were incubated in PBS/4% BSA/0.1% Triton X-100 with a primary antibodies overnight at 4°C, washed in PBS, incubated for 1 h with secondary antibodies, thoroughly washed in PBS, incubated with 4′,6′-diamidino-2-phenylindole (DAPI) for nuclear staining for 5 min. For the quantification of necrosis, sections were incubated only with a secondary anti-mouse antibody to reveal the absorption of immunoglobulin G by muscle necrotic fibers. Sections were then mounted in Fluoromount (Southern Biotech) and cells were maintained in PBS.

For PAX7 immunostaining, cryosections were rehydrated in PBS, fixed with PFA 4% for 10 min, permeabilized with precooled Methanol for 6 min and antigens were unmasked in citrate solution (Vector laboratories) at 95°C for 15 min. Cryosections were then blocked in PBS/5% BSA/10% HS/ 0.2% Triton X-100 for 1h then incubated with a primary antibodies overnight at 4°C, washed in PBS, incubated for 1 h with secondary antibodies, thoroughly washed in PBS, incubated with DAPI for nuclear staining for 5 min and mounted in Fluoromount (Southern Biotech).

Imaging was performed on a confocal Nikon Ti2 microscope equipped with a motorized stage. Antibodies used are listed in table S2.

### Morphometric analysis

For fiber and nucleus detection, we used different artificial intelligence deep learning pretrained models. Fibers were segmented with Cellpose v3^57^ using a model trained for cytoplasm. Nuclei were segmented using StarDist^58^. The segmented image masks were then imported into QuPath^59^, where we measured morphological features such as area and shape, and identified cells using the pixel intensity thresholding function.

CD45-positive signal was quantified on tissue sections using Fiji (ImageJ) by applying a threshold to isolate positive staining, followed by binarization and measurement of the positive area using the "Analyze Particles" function. The CD45-positive area (in µm²) was then normalized to the total tissue area analyzed to obtain the signal density per µm².

Finally, all measurements were exported to Excel, where we organized the data and performed statistical analyses such as CSA, fiber type classification, and staining positivity using thresholds.

### Immunoblotting

Cryosections from frozen TA muscles were homogenized with a Dounce homogenizer in a cell lysis buffer (Cell Signaling) supplemented with phosphatase inhibitor cocktail (Roche). Lysates were centrifuged for 5 min at 1500g and protein concentration was determined in supernatant with Bradford method using Protein Assay Dye Reagent (Bio-Rad). Proteins were denatured at 95°C for 5 min with Laemmli buffer and then separated by electrophoresis (Nu-PAGE 4–12% Bis-Tris gel; Thermofisher Scientific) and transferred to nitrocellulose membranes (GE Healthcare). Membranes were blocked with 5% skimmed milk diluted in Tris Buffered Saline (TBS)-0.1% Tween (TBS-T) and incubated at 4°C overnight with primary antibodies. After washes in TBS-T, membranes were incubated with secondary antibodies conjugated to a fluorophore (Bio-Rad). Images were acquired with ChemiDoc™ MP (Bio-Rad) and band intensity was quantified using Image Lab software (Bio-Rad). Antibodies used are listed in table S2.

### snRNAseq

#### Library preparation and sequencing

Tibialis anterior muscles were isolated from mice immediately following euthanasia, minced, snap frozen in liquid nitrogen and cryoconserved at - 80°C. Nuclei from n=3 (mdx and mdx-GDF5) and n=2 (WT) were isolated following Chromium Nuclei Isolation Kits (10x Genomics) according to the manufacturer’s instruction. For counting, nuclei were labeled with Acridine Orange and Propidium Iodide and counted using a Luna FL automatic cell counter. Nuclei from each sample were loaded onto separate lanes of the 10x Genomics Chromium with the aim to recover 20,000 nuclei per sample, using loading volumes recommended by the manufacturer. We employed Chromium Next GEM Single Cell 3′ Kit version 3.1 (10x Genomics). Libraries were generated according to the manufacturer’s instructions. Sequencing was performed on a NextSeq2000 (Illumina) targeting 100 million reads per nucleus.

#### Analysis

Fastq sequence were were loaded in R (version 4.4.1) and pre-processed using SoupX^60^. Sequencing data were demultiplexed and mapped with Cell Ranger software (10x Genomics). Mouse mm10 (Ensembl) reference was used for the alignment. Cell Ranger outputs (filtered_feature_bc_matrix) pre-processed and analyzed using Seurat version 5^61^. Each sample was pre-filtered as follows: nuclei with more than 5% mitochondrial, less than 200 expressed transcripts were excluded. As well as nuclei with more than 4500 expressed transcripts. Clustering analysis was perform using^32^ makers, to which BMP6 for the quiescent satellites^62^ and Hmga2 for the activated satellites^33^ were added. For differentially expressed genes (DEGs) were calculated with fold-change and p-value between mdx-GDF5 and mdx groups. GO analysis were performed using DEenrichRPlot function of the Seurat V5 R package with the "GO_Molecular_Function_2023’’ database from EnrichR database^63^ with the following thresholds: logfc.threshold = 0.25, max genes = 150. To identify and visualize the cell-cell communication changes between mdx-GDF5 and mdx cells, the CellChat V2^64^ R package was used according to developer tutorial (https://github.com/sqjin/CellChat). To visualize the changes in ligand-receptor expression between the mdx-GDF5 and mdx groups on quiescent or activated satellite cell populations, the function “netVisual_Chord_Cell” were used specifically for NCAM and VCAM pathways.

### RNA isolation and gene expression analysis

Total RNA was isolated from mouse muscle cryosections using TRizol (ThermoFischer Scientific) and Direct-zol RNA Miniprep (Zymo research) and from cells using NucleoSpin RNA II (Macherey-Nagel, Hoerdt, France).

Complementary DNA was generated with Superscript II Reverse Transcriptase (ThermoFischer Scientific), and analyzed by real-time qPCR. Real-time qPCR was performed on QuantStudio 3 Pro Real-Time PCR System (Applied Biosystems) using Power SybrGreen PCR MasterMix (Applied Biosystems). All data were analyzed using the 2^-ΔΔCT^ method and normalized to RPLP0 (Ribosomal protein lateral stalk subunit PO) mRNA expression. The sample reference to calculate mRNA fold change is indicated in each panel. Primers used are listed in table S1.

### Vector genome quantification

Genomic DNA was extracted from mouse muscles using the Puregene tissue kit (Qiagen). AAV genome copy number and genomic DNA were determined on 40 ng of genomic DNA by absolute quantitative real-time PCR on QuantStudio 3 Pro Real-Time PCR System (Applied Biosystems) with TaqMan™ Fast Advanced Master Mix (Applied Biosystems). The primers (forward: 5′-CATCAATGGGCGTGGATAGC-3′ and reverse: 5′-GGAGTTGTTACGACATTTTGGAAA-3′) and probe (5′-ATTTCCAAGTCTCCACCC-3′) were selected for specific amplification of the vector genome sequence. As a reference sample, a pAAV plasmid was 10-fold serially diluted (from 10^8^ to 10^2^ copies). All genomic DNA samples were analysed in duplicates.

### ELISA for Anti-AAV10 Antibody Detection

96-well ELISA plates (Nunc MaxiSorp™) were coated overnight at 4°C with 50 µL/well of AAV10 capsid diluted in carbonate buffer (5.10^8^/well). Plates were washed three times with PBS + 0. 5% Tween-20 (PBS-T) and blocked for 1 h at room temperature with 200 µL/well of 2% BSA in PBS-T.

Sera were diluted in blocking buffer (1:4000) and added to the plates (50 µL/well) in duplicate. Incubation was performed for 1 h at room temperature.

After washing, plates were incubated with 100µl of HRP-conjugated anti-mouse IgG (1:10000 dilution in blocking buffer) for 1 h at room temperature. After a final wash, 50 µL of TMB substrate was added per well, and the reaction was stopped after 10 min with 50 µL of 3M H₂SO₄.

Absorbance was measured at 450 nm using a microplate reader (e.g., BioTek or Thermo Scientific).

Negative controls (sera from non-injected mice) and positive controls (known anti-AAV10 positive sera) were included for comparison.

### Statistics analyses

For comparison between two groups, data were tested for normality using a Shapiro–Wilk test and for homoscedasticity using a Bartlett test followed by parametric (two-tailed paired, unpaired Student’s t-tests) or non-parametric test (Mann-Whitney) to calculate P values (as detailed in the figure legends). For comparison among more than two groups, according to normality ordinary one-way ANOVA or Kruskal-Wallis tests were performed. According to homoscedasticity, Brown-Forsythe ANOVA were performed. When it was necessary, two-way ANOVA tests were performed to compare between groups (as detailed in the figure legends). All ANOVA and Kruskal-Wallis were followed by appropriated post-hoc tests. All statistical analyses were performed with GraphPad Prism 8 software. Statistical significance was set at P < 0.05 and all bar graphs presented are means ± SEM. Asterisks indicate significant differences (*P < 0.05; **P < 0.01; ***P < 0.001) between groups according to statistical analysis performed.

## Acknowledgements

We acknowledge UMS28 animal facility for animal care and technical support and GENOM’IC platform at Cochin Institut for performing the single-nuclei RNA sequencing experiments.

## Author contributions

CG, AB and FPR conceived and designed the study. CG, AB, AV, BC, ZG, LS, PM and BHC performed the experiments. CG, AB, AV and LG analyzed the data. CG, AB and FPR wrote original draft and LG and SF reviewed the manuscript. FPR supervised the project and obtained the funding, as principal investigator, from the French National Research Agency (ANR) with LG as collaborator.

## Materiel & Correspondence

Requests should be addressed to

## Legends

**Figure S1.**
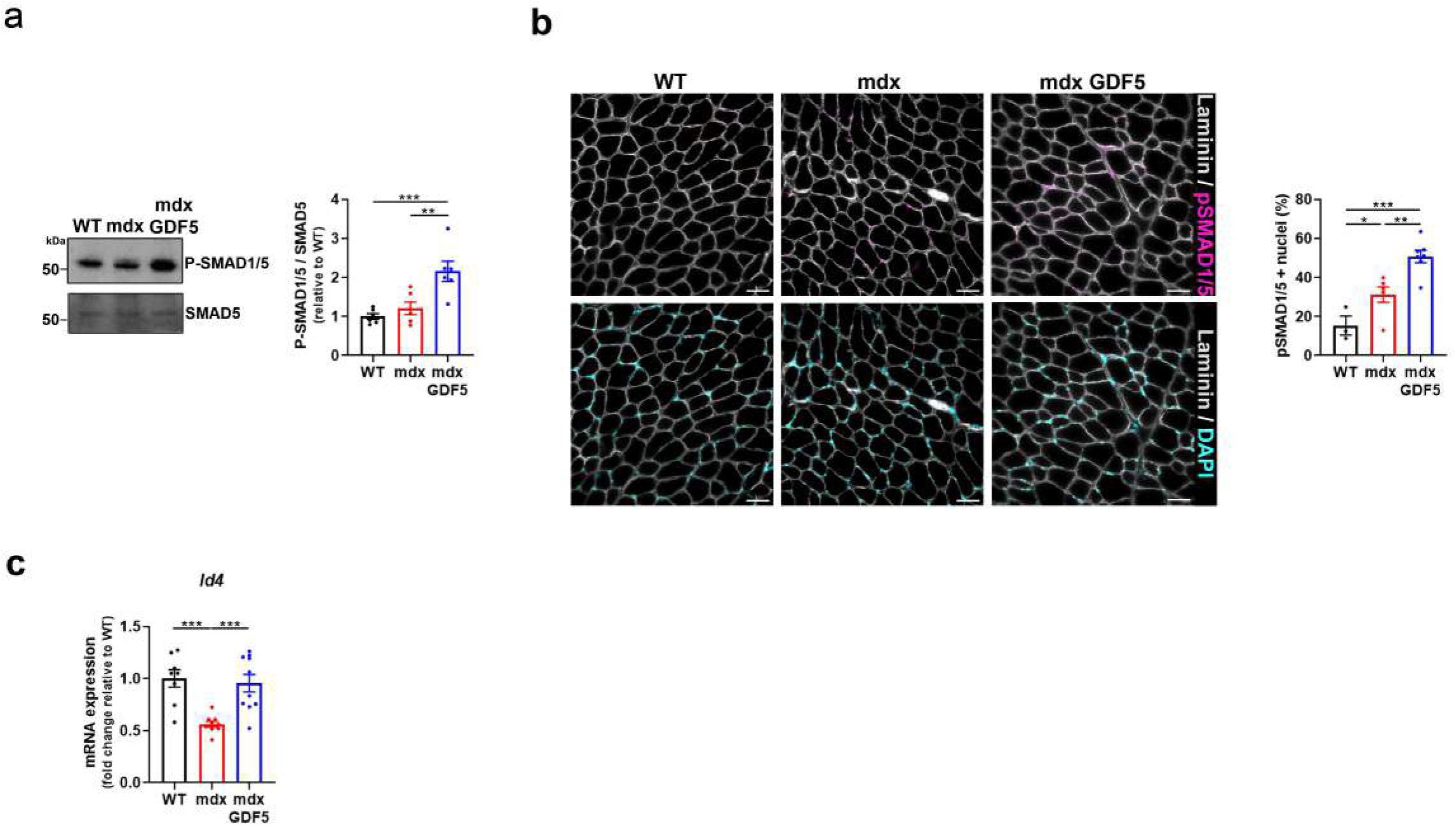
GDF5-OE activates SMAD1/5 pathway. **a**: Representative Western blot of TA lysates from WT, mdx and mdx-GDF5, 2 p.i., probed for P- SMAD1/5 and SMAD5. P-SMAD1/5 levels quantified and normalized to SMAD5 (n=6). **b**: Representative images of TA cryosections from WT, mdx and mdx-GDF5 2p.i. immunostained with P-SMAD1/5 (purple), laminin (grey) and DAPI (cyan), scale bar: 50µm. Percentage of P-SMAD1/5 positive nuclei. (n=3-7). **c**: RT-qPCR of *Id4* expression in TA from from WT, mdx, and mdx-GDF5, 2 p.i. (n=8-10). Data are presented as means ± s.e.m. *p-values* were calculated by Ordinary One-Way ANOVA test followed by Uncorrected Fisher LSD (a, b); Brown Forsythe One-Way ANOVA test followed by Unpaired t with Welch’s correction (c).

**Figure S2:**
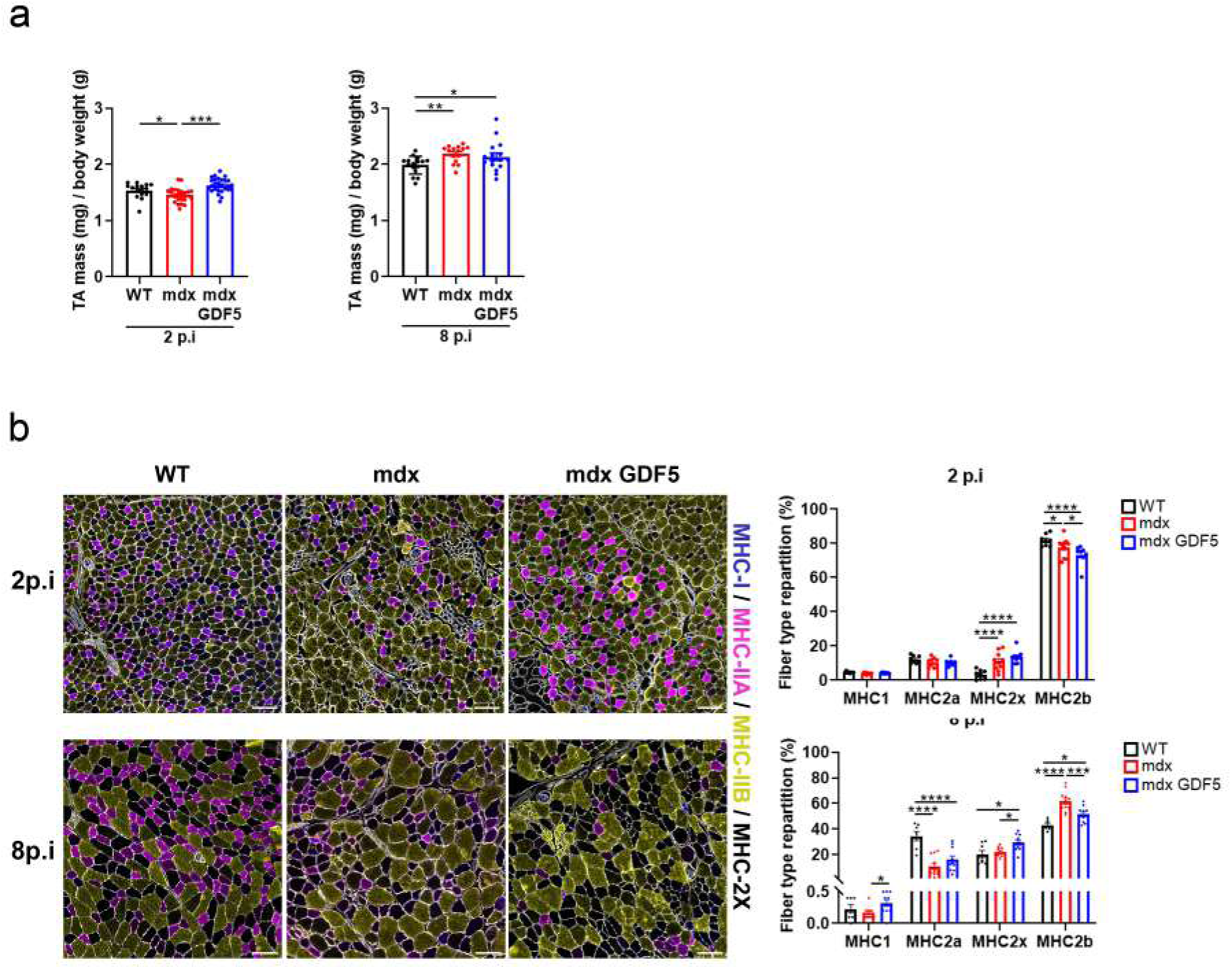
GDF5 modulates muscle mass and fiber type composition. **a:** TA mass relative to body weight. (2 p.i.: n=16-28); (8 p.i.: n=15-16). **b:** Representative image of cryosection from WT, mdx and mdx-GDF5 2- and 8-p.i. immunostained with laminin (grey), MHC-I (blue), MHC-IIA (purple), MHC-IIB (yellow) and MHC-IIX (black). scale bar 100 μm. Distribution of MHC fiber type (2 p.i.: n=8-10) (8 p.i.: n=7-11). Data are presented as means ± s.e.m. *p-values* were calculated Kruskal-Wallis test followed by Uncorrected Dunn’s test (a); Ordinary One-Way ANOVA test followed by Uncorrected Fisher LSD (a); Two-Way ANOVA test followed by Tukey’s multiple comparisons test (b).

**Figure S3:**
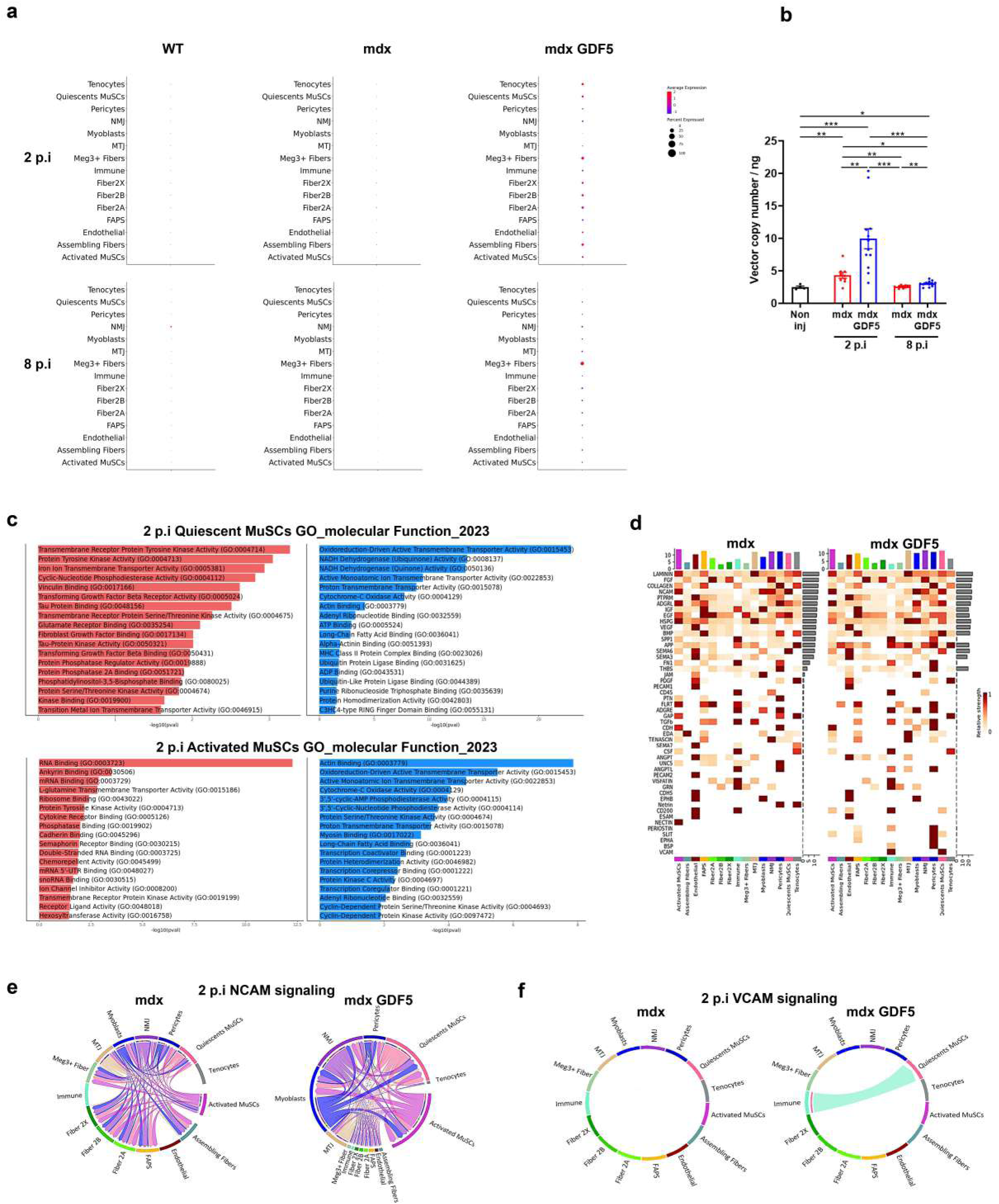
*Gdf5* transcript, AAV vector expression and signalling pathways in dystrophic muscles. **a:** Dotplots assessing Gdf5 expression for each major cell type in different conditions: WT, mdx and mdx-GDF5 2- and 8-p.i.. **b:** AAV genome copy quantification was determined in TA from mdx and mdx-GDF5 2- and 8-p.i. and compared to non-injected mdx muscles. Data are presented as means ± s.e.m. *p-values* were calculated by Brown Forsythe One-Way ANOVA test followed by Unpaired t with Welch’s correction. **c**: Top 18 upregulated (red) and downregulated (blue) Gene Ontology (GO) terms 2 p.i. in mdx compared to WT in Quiescents MuSCs (up) and Activated MuSCs (down) **d**: Heatmap showing the summary of overall signalling in mdx and mdx-GDF5. The color bar represents the relative signalling strength of a signalling pathway across cell types. The bars indicate the sum of the signalling strength of each cell type or pathway. **e,f**: Circle plot of pathway network for (e) NCAM and (f) VCAM pathways between mdx and mdx-GDF5 2 p.i..

**Figure S4:**
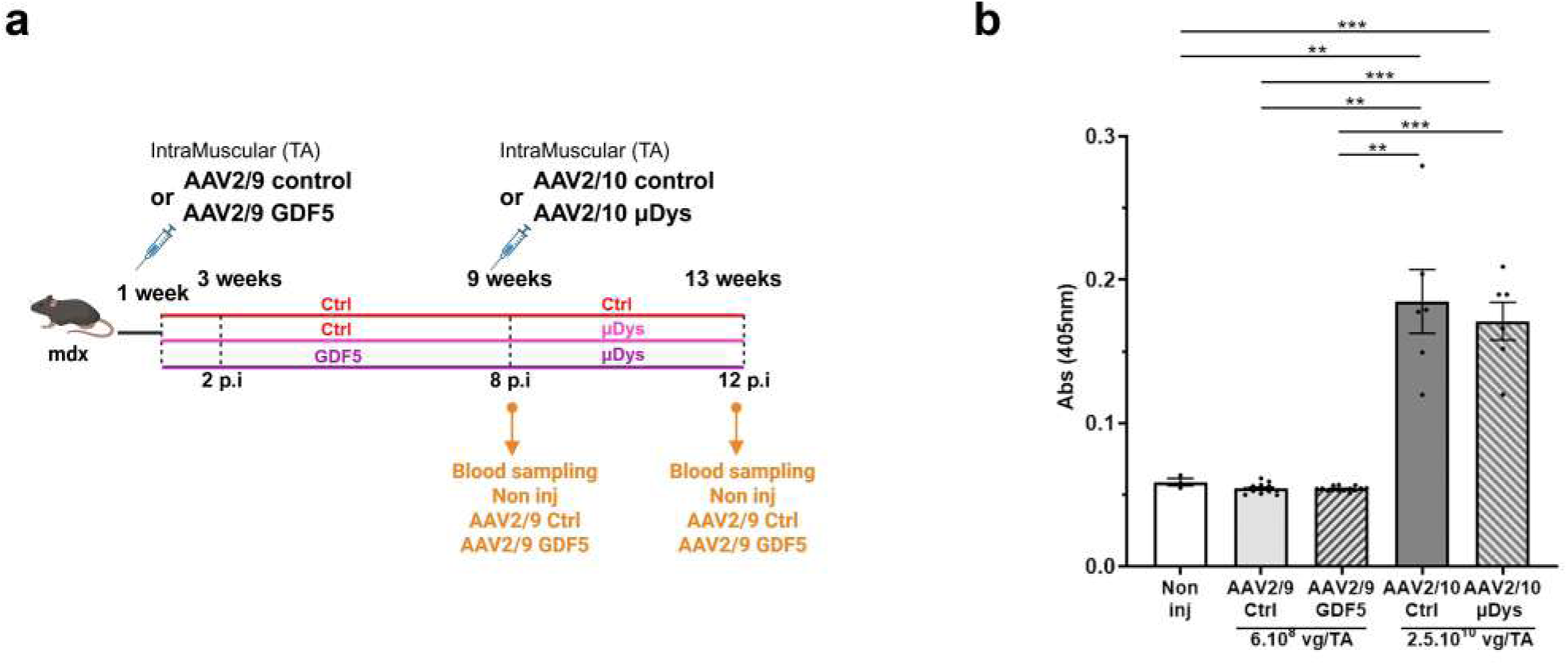
AAV2/9 injection does not induce production of anti-AAV2/10 antibodies. Anti-AAV10 antibody levels were measured in mdx TA injected with AAV2/9-control or -GDF5 at the dose of 6.10^8^ vg/TA during 8 weeks to assess cross-reactive immune responses. Serum samples from mdx TA injected with AAV2/10-control or -µDys at the dose of 2,5.10^10^ vg/TA were used as positive control. Serum from non-injected mice were used as negative control. Data are presented as means ± s.e.m. *p-values* were calculated by Brown Forsythe One-Way ANOVA test followed by Unpaired t with Welch’s correction.

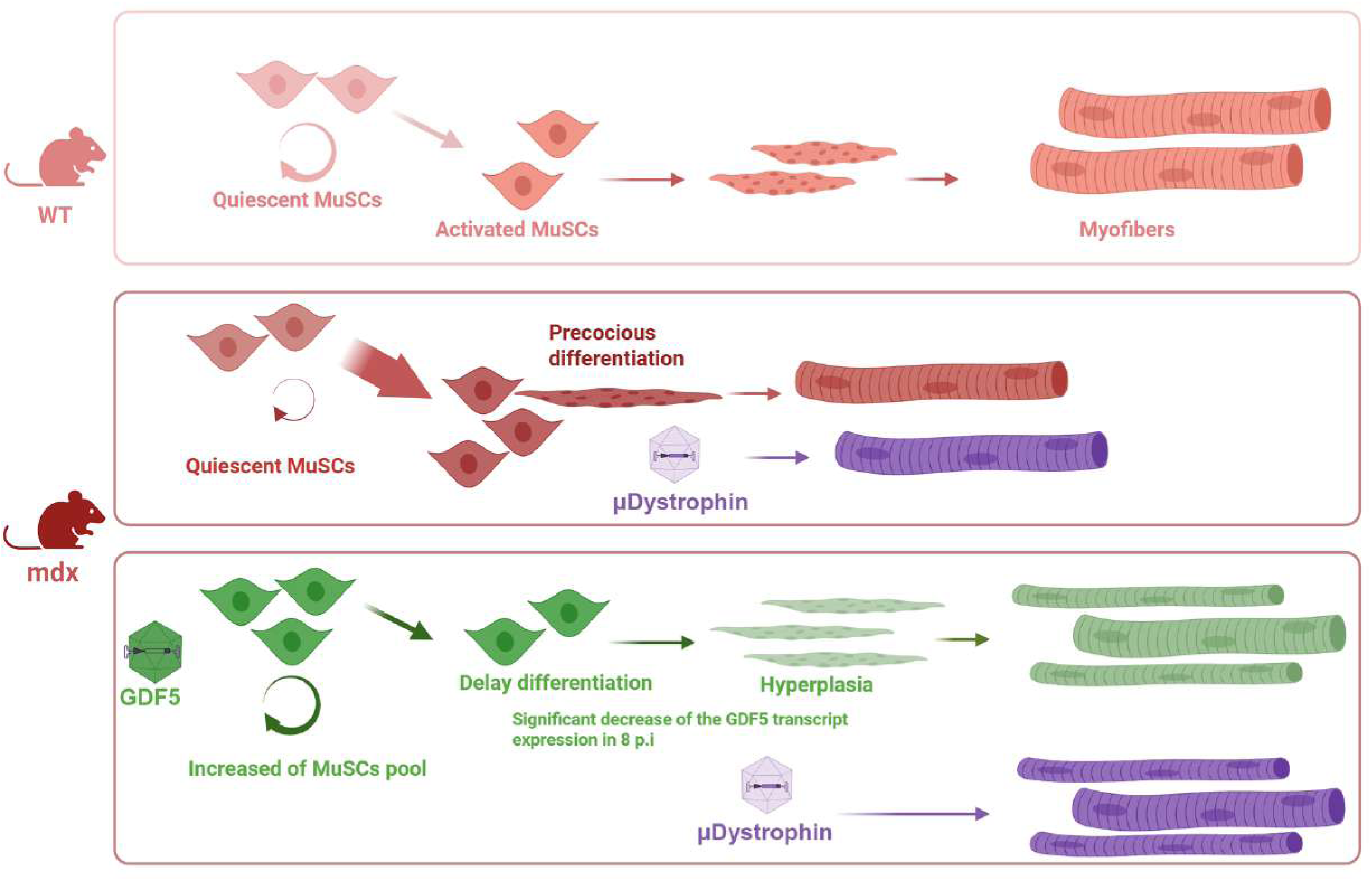

**Table S1:**
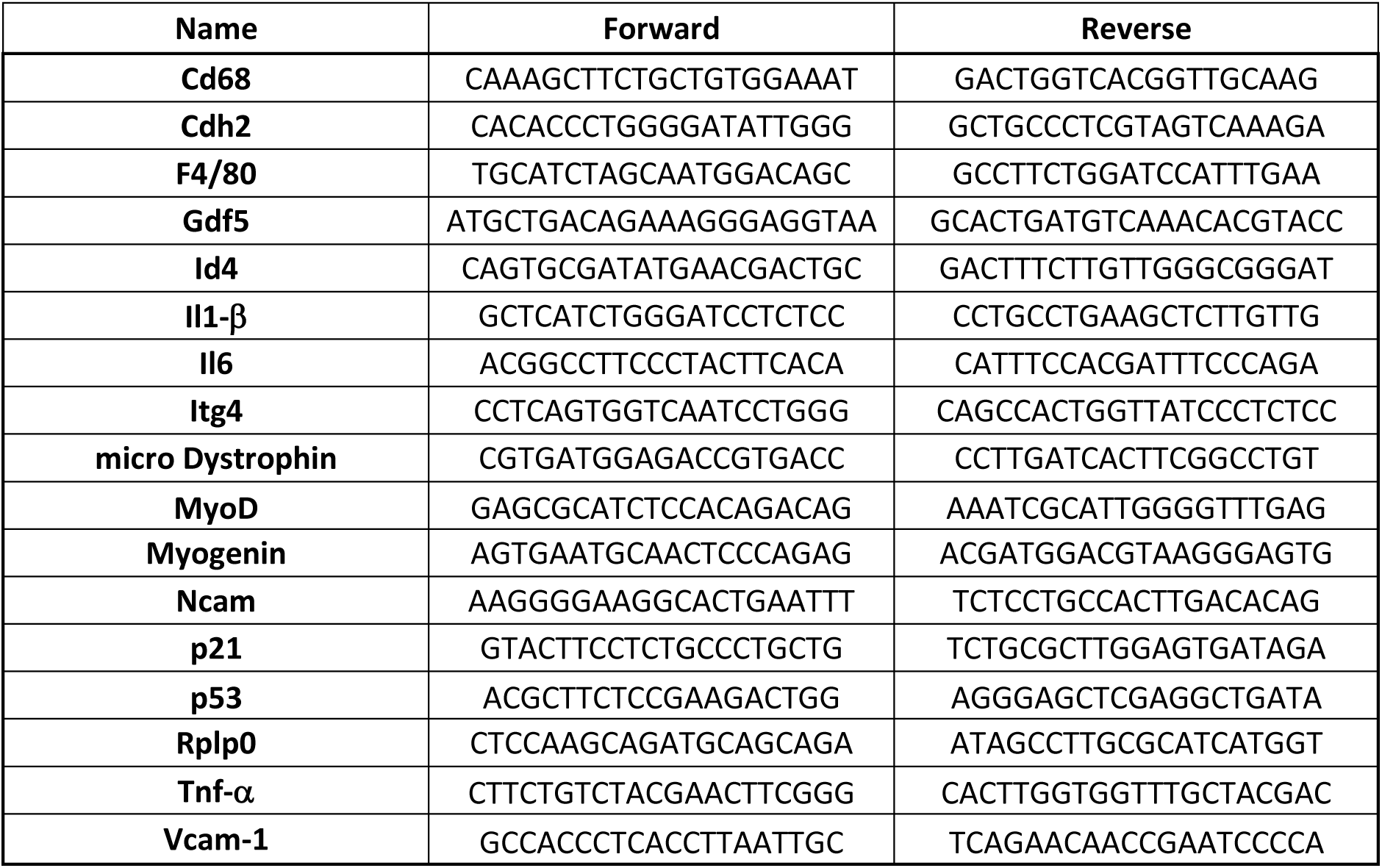
Table of primers used for quantitative PCR.

**Table S2:** Table of antibodies used for Immunofluorescence, Immunoblotting and FACS.

| Name / Target | Source | Reference | Species | Application | Dilution |
| --- | --- | --- | --- | --- | --- |
| Desmin | Proteintech | 16520-1-AP | Rabbit | IF | 1/500 |
| Ki67 | Abcam | ab15580 | Rabbit | IF | 1/500 |
| GDF5 | Santa Cruz | sc-373744 | mouse | WB | 1/500 |
| Laminin | Merck Millipore | L9393 | Rabbit | IF | 1/500 |
| MF-20 | DSHB | MF20-s | Mouse | IF | 1/20 |
| MYHI | DSHB | BA-D5-s | Mouse | IF | 1/10 |
| MYHIIa | DSHB | SC-71s | Mouse | IF | 1/10 |
| MYHIIb | DSHB | BF-F3-s | Mouse | IF | 1/10 |
| MYH3 | Santa Cruz | sc-53091 | Mouse | IF | 1/100 |
| PAX7 | Santa Cruz | sc-81648 | Mouse | IF | 1/50 |
| p-SMAD 1/5 | Cell signaling | 9516S | Rabbit | IF ; WB | 1/500 |
| CD31 eFluor 450 | eBioscience | 48-0311-82 | Mouse | FACS | 1/600 |
| CD45 eFluor 450 | eBioscience | 48-0451-82 | Mouse | FACS | 1/600 |
| Ly-6A/E (Sca-1) FITC | eBioscience | 11-5981-82 | Mouse | FACS | 1/600 |
| CD106 (VCAM-1) | eBioscience | 12-1061-82 | Mouse | FACS | 1/2000 |
| α7 Integrin | AbLab | R2F2 | rat | FACS | 1/300 |

